# Single-nucleus transcriptomics identifies a shared vulnerable excitatory neuronal population across typical and atypical Alzheimer’s disease

**DOI:** 10.64898/2026.03.30.715299

**Authors:** Felipe L. Pereira, Caroline Lew, Song H. Li, Liara Rizzi, Alexander V. Soloviev, Vitor Paes, Sara D. Brooks, Salvatore Spina, Jessica E. Rexach, Kathy L. Newell, Renata E. P. Leite, William W. Seeley, Claudia K. Suemoto, Bernardino Ghetti, Melissa E. Murray, Lea T. Grinberg

**Affiliations:** Department of Neuroscience, Mayo Clinic, Jacksonville, Florida, USA; Department of Quantitative Health Sciences, Mayo Clinic, Jacksonville, Florida, USA; Fein Memory and Aging Center, Weill Institute for Neurosciences, University of California San Francisco, San Francisco, USA; Physiopathology in Aging Laboratory (LIM-22), Department of Internal Medicine, University of Sao Paulo Medical School, Sao Paulo, Brazil; Program in Neurogenetics, Department of Neurology, David Geffen School of Medicine, University of California Los Angeles, Los Angeles, USA; Department of Human Genetics, David Geffen School of Medicine, University of California Los Angeles, Los Angeles, USA; Department of Pathology and Laboratory Medicine, Indiana University School of Medicine, Indianapolis, USA; Department of Pathology, University of Sao Paulo Medical School, Sao Paulo, Brazil; Division of Geriatrics, Department of Internal Medicine, University of Sao Paulo Medical School, Sao Paulo, Brazil

**Keywords:** atypical AD, neuronal selective vulnerability, lvPPA, PCA

## Abstract

Alzheimer’s disease (AD) presents with substantial clinical and anatomical heterogeneity, including both typical amnestic and atypical variants such as posterior cortical atrophy and logopenic primary progressive aphasia. Although neurofibrillary tangle (NFT) burden is a defining pathological feature of AD, its regional distribution varies across clinical phenotypes, suggesting that selective neuronal vulnerability may shape disease presentation. However, the cellular and molecular determinants underlying this vulnerability remain incompletely understood. Here, we profiled single-nucleus transcriptomes across multiple brain regions, including hippocampal (CA1) and neocortical (superior temporal gyrus and occipital cortex) regions, from individuals with typical and atypical AD and healthy controls. Integrative analysis identified major cell classes and resolved diverse excitatory and inhibitory neuronal subpopulations that were reproducibly observed across regions and individuals. Using quasi-binomial regression models to assess compositional changes, we quantified subtype-specific vulnerability associated with AD pathology. We identified a distinct excitatory neuronal subpopulation characterized by NRGN and BEX1 expression, which showed reproducible depletion across multiple regions, with the strongest evidence in amnestic AD and in neocortical regions in lvPPA. This vulnerable population showed concordance with previously reported AD-associated excitatory neuron signatures, supporting a conserved transcriptional program of susceptibility. Genes enriched in this population were associated with chemical synaptic transmission and regulation of synaptic plasticity and formed interconnected networks in protein-protein interaction analyses. These findings suggest that intrinsic properties related to synaptic function may predispose specific neuronal populations to degeneration in AD. Together, our results define a conserved, transcriptionally distinct excitatory neuron subpopulation that is selectively vulnerable across AD phenotypes and brain regions. This work provides a framework for linking regional pathology to cell-type-specific susceptibility and highlights synaptic regulatory pathways as potential contributors to neuronal degeneration in Alzheimer’s disease.

## Introduction

Alzheimer’s disease (AD) most commonly presents as a progressive amnestic syndrome, but a substantial subset of cases, particularly among younger-onset patients, manifests atypical clinical phenotypes dominated by language, visuospatial, executive, behavioral or motor dysfunction [1–3]. Among these atypical phenotypes, the logopenic variant of primary progressive aphasia (lvPPA) and posterior cortical atrophy (PCA) are especially informative because they couple a shared AD neuropathological substrate to sharply different cognitive syndromes and neuroanatomical signatures [2–5]. This clinical diversity suggests that the same core disease process can engage partially distinct neural systems, raising the possibility that selective neuronal vulnerability is a major determinant of AD phenotype.

This heterogeneity is mirrored at the neuropathological level. Although Braak staging captures the stereotyped progression of tau pathology through limbic and neocortical territories, AD also includes clinicopathological subtypes with distinct regional distributions of neurofibrillary tangles (NFTs) [6]. In cohorts enriched for atypical presentations, cortical—but not hippocampal—NFT burden is greater in atypical variants than in amnestic AD. Prior clinicopathological work further showed that lvPPA has disproportionate NFT accumulation in the superior temporal gyrus (STG), whereas PCA shows disproportionate involvement of the occipital cortex (OCP); Cornu Ammonis sector 1 (CA1) of the hippocampus is also heavily affected, particularly in amnestic AD [7]. These observations indicate that NFT burden is not simply a scalar measure of disease severity, but rather follows regionally biased patterns that map onto clinical phenotype. Typical and atypical AD therefore provide a natural framework for testing whether vulnerable neuronal populations are shared across syndromes or instead track phenotype-linked regional pathology.

Single-cell and single-nucleus transcriptomic approaches now allow direct interrogation of selective vulnerability in human brain tissue. Prior studies have shown that AD progression is accompanied by reproducible changes in cellular composition and by depletion of specific excitatory neuronal populations with distinctive molecular identities [8–10]. In particular, our group identified selectively vulnerable excitatory neuron subpopulations in the entorhinal cortex and superior frontal gyrus [8], whereas Mathys and colleagues demonstrated the power of single-cell transcriptomics to resolve disease-associated cellular states in the neocortex [9] and later extended this approach to a multiregional framework that uncovered vulnerable neuronal populations across six brain regions [10]. Together, these studies show that neuronal vulnerability in AD is molecularly structured rather than random. However, they did not directly leverage neuropathologically matched typical and atypical AD phenotypes to test whether the neuronal states most vulnerable in AD represent a shared disease-wide substrate or differ according to syndrome-linked regional tau topography.

Here, we hypothesized that typical and atypical AD would converge on a shared vulnerable excitatory neuronal population despite differences in regional NFT distribution, but that the molecular context surrounding this vulnerable state would vary by phenotype and brain region. To test this, we applied single-nucleus RNA sequencing (snRNA-seq) to three selectively vulnerable regions, hippocampal CA1, STG, and OCP, from neuropathologically confirmed amnestic AD, lvPPA, PCA, and healthy control (HC) brains. We then asked whether vulnerable neuronal populations were shared or syndrome-enriched across these phenotypes, whether excitatory and inhibitory neuronal compartments were affected similarly, and whether the most vulnerable neuronal population carried a conserved molecular program that might help explain its susceptibility to tau-associated degeneration.

## Material and Methods

### Human cohort and regional sampling

Post-mortem human brain tissue was obtained from the Brain Bank of the Department of Neuroscience at Mayo Clinic, the Neurodegenerative Disease Brain Bank at the University of California, San Francisco, the Biobank for Aging Studies at the University of São Paulo, and the Brain Bank at Indiana University School of Medicine. The study cohort comprised 68 individuals spanning four groups: HC (n = 27), amnestic AD (n = 24), lvPPA (n = 11), and PCA (n = 6) (Table 1; case-level metadata in Supplementary Table 1). Across all donors, 168 donor-region samples were profiled, including CA1 (n = 51), STG (n = 60), and OCP (n = 57). These three regions were selected a priori on the basis of prior clinicopathological work showing preferential CA1 involvement in amnestic AD, STG involvement in lvPPA, and OCP involvement in PCA [7].

**Table 1.**
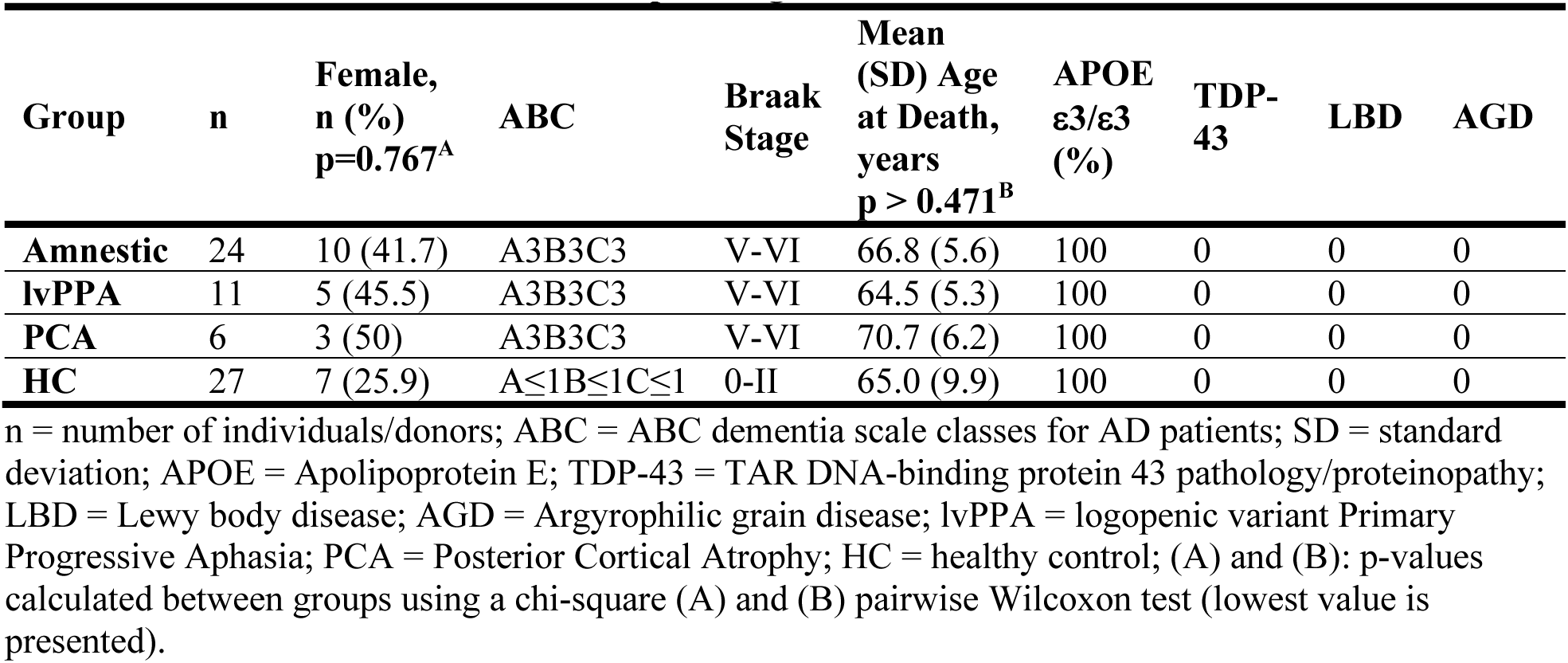
Cohort characteristics and neuropathological features p=0.767.

Clinical diagnoses were assigned using established consensus criteria for primary progressive aphasia and posterior cortical atrophy and were subsequently confirmed by post-mortem neuropathological evaluation [4,5]. Neuropathologic assessment followed National Institute on Aging–Alzheimer’s Association recommendations [11]. All AD cases met high-level AD neuropathologic change (ABC score A3B3C3) and Braak NFT stages V-VI. HC cases showed low or absent AD neuropathologic change (ABC ≤ A1B1C1) and Braak stages 0-II. To minimize genetic and pathological heterogeneity, all individuals were APOE ε3/ε3 carriers, and cases with comorbid TDP-43 proteinopathy, Lewy body disease, or argyrophilic grain disease were excluded. All procedures were performed under approval from the relevant institutional review boards at the participating sites.

### Isolation of nuclei from frozen human brain tissue

Frozen tissue was processed using detergent-based Dounce homogenization followed by OptiPrep density-gradient purification. All steps were performed on ice or at 4 °C. Briefly, tissue was transferred to chilled sucrose-based homogenization buffer containing divalent cations, reducing agent, polyamines, protease inhibitors and RNase inhibitor, and homogenized with a glass Dounce. IGEPAL CA-630 was then added to lyse cellular membranes while preserving nuclear integrity, followed by additional Dounce homogenization. Homogenates were filtered through a 40 µm strainer, mixed with OptiPrep working solution, layered onto a discontinuous 30%/40% OptiPrep gradient and centrifuged at 10,000g at 4 °C. Nuclei were collected from the 30%/40% interphase, washed in PBS containing 0.04% BSA, counted and diluted to the target concentration for droplet-based library preparation.

### snRNA-seq library preparation and sequencing

Single-nucleus 3′ gene-expression libraries were generated with Chromium Single Cell 3′ Reagent Kits v3 (10x Genomics) following the manufacturer’s protocol without modification. Purified nuclei were loaded onto the Chromium Controller to generate gel bead-in-emulsions,followed by reverse transcription, cDNA amplification and library construction. Final libraries were quality-controlled and sequenced on an Illumina NovaSeq instrument using standard 10x Genomics run parameters.

### Read processing, quality control and integration

Raw sequencing data were processed with Cell Ranger v9.0.0 (10x Genomics) using a GRCh38 pre-mRNA reference to capture nuclear transcripts. Ambient/background RNA was removed with CellBender remove-background v0.3.1 [12]. Corrected matrices were imported into Seurat v5 [13]. Nuclei expressing fewer than 200 genes, or with >5% mitochondrial RNA or >5% ribosomal RNA, were excluded. Data were normalized with SCTransform [14], and doublets were identified independently for each donor-region sample with scDblFinder v1.22 [15].

Filtering thresholds were applied uniformly across samples. Clusters showing residual doublet signatures or strong sample-specific artefacts were excluded during manual curation. The final curated dataset comprised 1,615,604 nuclei.

Initial integration, clustering, and lineage assignment were performed without using diagnostic labels. Tissue source and postmortem interval (PMI) were evaluated during exploratory analyses as potential confounders and did not materially alter clustering structure or the direction of donor-level downstream results; they were therefore not included as primary covariates in the final models. For clustering construction, 150,000 nuclei were selected using the SketchAssay framework in Seurat v5 [13]. Principal component analysis was performed on the 4,000 most variable genes, Harmony [16] was used for sample alignment, and Leiden clustering was performed on the shared nearest-neighbor graph at a resolution of 2.0. Cluster identities were then projected back onto the full dataset.

### Cell-type and subtype annotation

Major cell classes were annotated in Seurat using module scores derived from curated marker sets and canonical lineage markers. Excitatory neuronal identities were guided by NRGN, CUX2, RORB, THEMIS, TLE4 and ERBB4, whereas inhibitory neuronal identities were guided by GAD1, VIP, LAMP5, SST and PVALB. Non-neuronal populations were identified using established markers for astrocytes, microglia, oligodendrocytes, oligodendrocyte precursor cells (OPCs), endothelial cells, vascular leptomeningeal cells (VLMCs), T cells and fibroblasts [26].

Excitatory and inhibitory neurons were then subsetted and reprocessed independently using the same workflow to resolve subtype-level structure. Marker genes were identified with FindAllMarkers using a Wilcoxon rank-sum test, requiring detection in at least 20% of nuclei within a cluster and a minimum log-fold change >0.10. To avoid circularity due to disease-related subtype naming, second-level neuronal subtype annotation was based solely on HC nuclei. To benchmark subtype assignments, we computed Spearman correlations between average expression profiles from our subtypes and published external reference atlases, restricting analyses to shared genes. Correlations were performed separately for excitatory, inhibitory and non-neuronal lineages.

### Modeling subtype vulnerability

Subtype-specific vulnerability was evaluated by modeling changes in the proportion of nuclei assigned to each neuronal subtype across diagnostic groups. Each analytical observation corresponded to one donor-region sample rather than to individual nuclei. For a given subtype, the observed proportion was represented as the number of nuclei assigned to that subtype (n.per.cluster) relative to the total number of nuclei in the corresponding parent class (total.per.sample). Subtypes not detected in a sample were assigned a count of zero.

For each subtype, we fit a generalized linear model with quasibinomial error and logit link: cbind(n.per.cluster, total.per.sample - n.per.cluster) ∼ PrimaryDx + Sex. Models were fit separately within each brain region (CA1, STG, and OCP). HC served as the reference level. Model coefficients were exponentiated to obtain odds ratios (ORs) and 95% confidence intervals (CIs), representing the odds that a nucleus belonged to a given subtype in each disease group relative to HC after adjustment for sex. P values were adjusted across subtypes within each neuronal lineage using the Benjamini–Hochberg false discovery rate (FDR) procedure. Because these models operate on cell-class composition, ORs quantify relative depletion or enrichment within the parent neuronal compartment rather than absolute cell counts.

### Pseudobulk differential expression and conserved gene enrichment

To identify molecular features associated with the vulnerable Exc NRGN BEX1 subtype, we restricted analyses to HC nuclei to minimize confounding by disease-associated transcriptional perturbation. For each brain region, raw counts were aggregated into donor-level pseudobulk profiles by subtype. Only regionally prevalent excitatory subtypes, defined as subtypes detected in at least 50% of samples from that region, were retained for comparison.

Within each region, DESeq2 [17] was used to compare Exc NRGN BEX1 against each other prevalent excitatory subtype in a pairwise fashion. Genes retained as enriched or depleted were required to show directional concordance across all pairwise contrasts within that region. Mean log2 fold change and mean adjusted P value across pairwise contrasts were used for ranking and visualization. Region-specific Exc NRGN BEX1 gene sets were then intersected across CA1, STG, and OCP to define a conserved BEX1-associated program for Fig. 4. Functional enrichment of the shared gene set was performed with STRING [18] using Gene Ontology biological-process terms and protein-protein interaction network analysis.

### Curated pathway-focused differential expression analysis

To assess whether the molecular context surrounding the shared vulnerable neuronal state differed across AD phenotypes, we performed a targeted differential-expression analysis using curated gene sets related to calcium dysregulation/neuroinflammation and mitochondrial dysfunction/oxidative stress. Pseudobulk count matrices were generated in Seurat by aggregating raw UMI counts across nuclei sharing the same Individual ID, brain region, and annotated cell cluster, yielding one pseudobulk profile per donor-region-cell-cluster combination. Differential expression was then performed using DESeq2 [17] within each region and cell cluster, comparing each AD phenotype with HC.

For visualization in Fig. 5, log2 fold-change estimates were row-standardized as Z-scores across comparisons and displayed as heatmaps. Black diamond symbols indicate gene-cell cluster comparisons meeting both statistical and effect-size thresholds (adjusted P < 0.05 and absolute log2 fold change > 0.5). Because the three AD phenotypes were unequally represented, these heatmaps were interpreted as descriptive, hypothesis-generating summaries of effect-size patterns rather than as formal between-syndrome tests.

### Multiplex immunofluorescence validation of the BEX1-positive excitatory neuronal population

For orthogonal validation in the superior temporal gyrus (STG), postmortem human brain tissue from healthy controls (HC, n = 5), amnestic AD (n = 5), lvPPA (n = 5), and PCA (n = 4) was analyzed by multiplex immunofluorescence. Formalin-fixed, paraffin-embedded [or frozen] STG sections from the same anatomical region were cut at []-µm thickness and processed in parallel to minimize batch effects. After deparaffinization and rehydration [if FFPE], sections underwent antigen retrieval in [buffer, pH, temperature, duration] and were treated with [autofluorescence-reduction step, if used]. Non-specific binding was blocked with [blocking solution], followed by incubation with primary antibodies against NeuN (neuronal marker), CAMKII/CaMKIIα (excitatory neuronal marker), BEX1 (population of interest), and PHF1 (phospho-tau). Primary antibodies, host species, vendors, catalog numbers, and working dilutions are listed in Supplementary Table. Appropriate species-specific secondary antibodies conjugated to spectrally distinct fluorophores were used, and nuclei were counterstained with [DAPI/Hoechst, if used].

Whole-slide fluorescent images were acquired on a Zeiss using identical acquisition settings across cases within each marker channel. For quantitative analysis, non-overlapping cortical fields per section were selected from gray matter in STG according to predefined anatomical criteria and excluding tissue folds, edge artifacts, large vessels, and areas with marked section damage. Field selection and quantification were performed blinded to diagnostic group. Images were analyzed in ImageJ/Fiji using a standardized pipeline that included channel separation, background subtraction, and thresholding held constant across cases. Cells were classified sequentially as NeuN-positive neurons, CAMKII-positive excitatory neurons within the NeuN-positive population, and BEX1-positive excitatory neurons within the CAMKII-positive neuronal population. PHF1 signal was used to visualize phospho-tau pathology in the same tissue sections but was not used to define the primary quantitative outcome. The primary outcome for Fig. 2e was the proportion of excitatory neurons that were BEX1-positive, calculated for each donor as: BEX1-positive excitatory neurons / total CAMKII-positive NeuN-positive neurons. Counts from multiple fields were first aggregated within donor to generate one donor-level summary value, and each donor contributed a single independent observation to the statistical analysis. Group differences were then tested across HC, amnestic AD, lvPPA, and PCA as described below.

### Statistical analysis

Demographic and neuropathological characteristics summarized in Table 1 were compared across groups using χ² tests for categorical variables and Wilcoxon rank-sum tests for age at death, as reported in Table 1. For STG histological validation, group differences were tested at the donor level using one-way analysis of variance followed by Tukey’s honest significant difference test.

Changes in the excitatory-to-inhibitory ratio were modeled using linear mixed-effects models with PrimaryDx, BrainRegion, Sex, and their interactions (PrimaryDx × BrainRegion and PrimaryDx × Sex) as fixed effects, and donor identity as a random intercept to account for repeated regional sampling from the same donor [19].

Unless otherwise specified, all tests were two-sided.

## Results

### An NFT-affected-region single-nucleus map of typical and atypical AD

To resolve cell-type-specific vulnerability across distinct AD phenotypes, we generated snRNA-seq data from hippocampal CA1, STG and OCP across 68 donors comprising amnestic AD (n = 24), lvPPA (n = 11), PCA (n = 6) and HC (n = 27) (Fig. 1a,b and Table 1). These regions were selected to capture the characteristic topography of NFT burden across AD variants, with CA1 predominantly affected in amnestic AD, STG in lvPPA, and OCP in PCA (Fig. 1a). All AD cases met high-level AD neuropathologic change (A3B3C3; Braak V-VI), whereas controls showed little or no AD pathology. Sex distribution and age at death were comparable across groups (Table 1).

**Figure 1.**
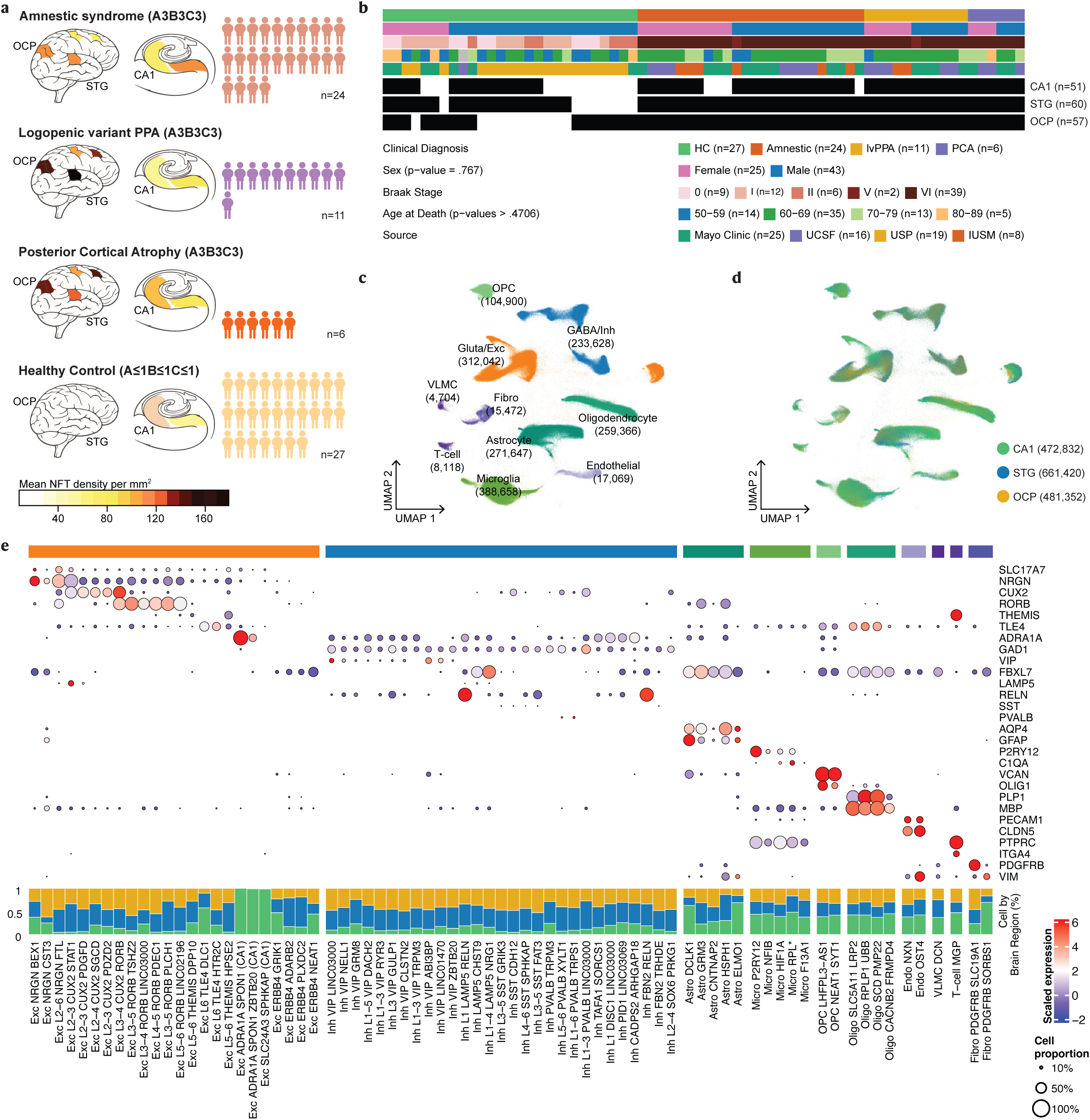
Cohort design and single-nucleus atlas across selectively vulnerable brain regions. a,b, Overview of the clinicopathological cohort, including diagnosis group, sampled brain regions, region-specific mean NFT density [7], demographic distributions, tissue sources and the number of donor-region samples profiled. c, UMAP of 1,615,604 nuclei colored by major cell-class annotation; class-specific counts are indicated. d, Same embedding colored by brain region (CA1, STG and OCP). e, Dot plot of canonical marker genes across neuronal and non-neuronal subclasses; dot size indicates the proportion of nuclei expressing each marker and color indicates scaled average expression.

After removal of low-quality nuclei, ambient RNA contamination and doublets, we retained 1,615,604 high-quality nuclei. These nuclei segregated into major excitatory and inhibitory neuronal, glial, vascular and immune classes (Fig. 1c), including excitatory and inhibitory neurons, astrocytes, microglia, oligodendrocytes, OPCs, endothelial cells, VLMCs, fibroblasts and T cells. Projection of all nuclei into a shared low-dimensional space showed that major cellular identities remained well separated across the full cohort (Fig. 1c). Coloring the same embedding by brain region revealed extensive intermixing of CA1, STG and OCP nuclei within each major lineage (Fig. 1d), indicating that cell identity was the dominant source of transcriptional structure at this level of resolution. Canonical marker expression confirmed the expected identities of neuronal and non-neuronal subclasses, including SLC17A7/NRGN/CUX2/RORB/THEMIS/TLE4 for excitatory neurons, GAD1/VIP/LAMP5/SST/PVALB for inhibitory neurons, AQP4/GFAP for astrocytes, P2RY12/C1QA for microglia, and PLP1/MBP for oligodendrocyte-lineage cells (Fig. 1e).

Reciprocal excitatory and inhibitory module scores further supported neuronal class assignment (Supplementary Fig. 2). Across diagnoses and regions, subtype-level profiles remained most similar within lineage, supporting stable annotation despite phenotypic heterogeneity (Supplementary Fig. 1). Correlation with external reference atlases also validated excitatory, inhibitory and non-neuronal subtype assignments, including CA1-restricted excitatory states, ERBB4-positive excitatory populations, canonical interneuron subclasses and glial/vascular populations (Supplementary Figs. 3-5).Given the stability of broad cell classes across regions, we next asked whether vulnerability emerged at the neuronal subtype level.

### A conserved NRGN-BEX1 excitatory population is selectively depleted across AD phenotypes

Sub clustering of glutamatergic neurons identified 24 excitatory subtypes organized along NRGN, CUX2, RORB, THEMIS/TLE4, CA1-restricted and ERBB4-positive axes (Fig. 2a). Several excitatory populations were broadly represented across all three sampled regions, including Exc NRGN BEX1 (94.1% of CA1 samples, 93.3% of STG samples and 72.4% of OCP samples), Exc L2-4 CUX2 SGCD (94.1%, 98.3% and 100.0%, respectively) and Exc L5-6 THEMIS DPP10 (68.6%, 70.0% and 65.5%) (Fig. 2b). By contrast, Exc L2-3 CUX2 PDGFD showed strong neocortical prevalence (54.9% in CA1, 100.0% in STG and 98.3% in OCP), whereas Exc ADRA1A SPON1 and Exc SLC24A3 SPHKAP were restricted to CA1.

**Figure 2.**
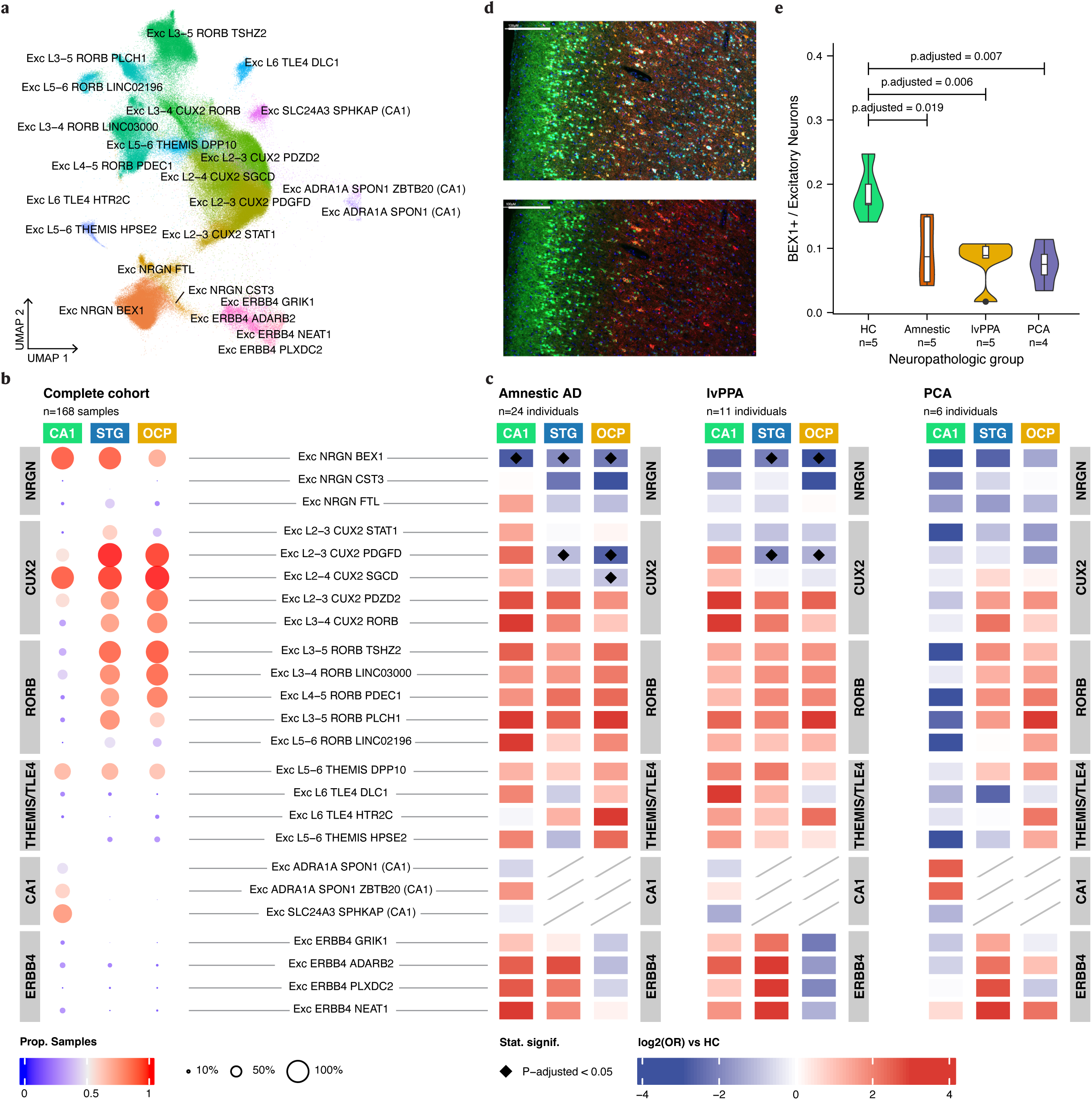
Conserved excitatory-neuron vulnerability across typical and atypical AD. a, UMAP of 24 excitatory neuronal subtypes. b, Proportion of region-specific samples in which each excitatory subtype was detected across the full cohort (n = 168 donor-region samples). c, Relative-abundance modeling of excitatory subtype vulnerability across diagnoses and regions. Values are log2-transformed ORs from quasibinomial models comparing each diagnosis with HC; blue indicates relative depletion and red indicates relative enrichment. Diamonds mark FDR-adjusted P < 0.05. Diagonal hatching depicts region-restricted/not-modeled. d, Representative multiplex STG images stained for NeuN, CAMKII, PHF1 and BEX1. Scale bars: 200 µm. e, Quantification of the proportion of BEX1-positive excitatory neurons in STG across diagnostic groups.

Relative-abundance modeling revealed that Exc NRGN BEX1 was the only excitatory population showing reproducible depletion across both typical and atypical AD phenotypes (Fig. 2c). In amnestic AD, Exc NRGN BEX1 was significantly depleted in CA1 (OR = 0.14, 95% CI 0.05-0.38, adjusted P = 0.021), STG (OR = 0.26, 95% CI 0.10-0.59, adjusted P = 0.013) and OCP (OR = 0.21, 95% CI 0.08-0.47, adjusted P = 0.003). Exc L2-3 CUX2 PDGFD was also depleted in STG (OR = 0.47, 95% CI 0.30-0.72, adjusted P = 0.006) and OCP (OR = 0.15, 95% CI 0.08-0.28, adjusted P = 1.1 × 10^-5^), whereas Exc L2-4 CUX2 SGCD showed region-restricted depletion in OCP (OR = 0.51, 95% CI 0.35-0.74, adjusted P = 0.004).

In lvPPA, Exc NRGN BEX1 again showed selective depletion in neocortex, reaching significance in STG (OR = 0.22, 95% CI 0.07-0.60, adjusted P = 0.027) and OCP (OR = 0.10, 95% CI 0.01-0.39, adjusted P = 0.025), but not in CA1 (OR = 0.19, 95% CI 0.06-0.56, adjusted P = 0.085). Exc L2-3 CUX2 PDGFD displayed a parallel pattern, with significant depletion in STG (OR = 0.28, 95% CI 0.15-0.49, adjusted P = 7.5 × 10^-4^) and OCP (OR = 0.41, 95% CI 0.19-0.81, adjusted P = 0.048). In PCA, no excitatory subtype showed FDR-significant depletion. Nevertheless, Exc NRGN BEX1 trended toward lower abundance in CA1 (OR = 0.07, 95% CI 0.001-0.50, adjusted P = 0.170) and STG (OR = 0.16, 95% CI 0.01-0.79, adjusted P = 0.130), and Exc L2-3 CUX2 PDGFD trended toward depletion in OCP (OR = 0.31, 95% CI 0.10-0.77, adjusted P = 0.057).

These depletions occurred alongside relative enrichment of several neocortical RORB-positive populations in PCA, indicating that excitatory compositional changes extended beyond the vulnerable states alone.

Because Exc NRGN BEX1 emerged as the only excitatory population depleted across amnestic AD and atypical neocortical phenotypes, we next sought orthogonal validation in STG tissue. Multiplex histology confirmed a significant reduction in BEX1-positive excitatory neurons across diagnostic groups (one-way ANOVA, F = 7.30, P = 0.00303; Fig. 2d,e). Post hoc Tukey comparisons showed that each AD syndrome had fewer BEX1-positive excitatory neurons than HC (adjusted P values ranging from 0.006 to 0.019), corroborating the compositional depletion observed in the single-nucleus data.

### Inhibitory-neuron compositional changes are more limited, but the excitatory-to-inhibitory balance is consistently reduced in AD

Sub clustering of GABAergic interneurons identified 29 inhibitory subtypes spanning canonical VIP, LAMP5, SST and PVALB classes, together with rarer interneuron states (Fig. 3a). In contrast to excitatory neurons, most inhibitory subtypes were broadly detected across the three sampled regions (Fig. 3b), including Inh VIP NELL1 (94.1% of CA1 samples, 100.0% of STG samples and 98.3% of OCP samples) and Inh L1-4 LAMP5 NRG1 (94.1%, 98.3% and 96.6%). Thus, inhibitory subtype architecture was largely conserved across hippocampal and neocortical territories.

**Figure 3.**
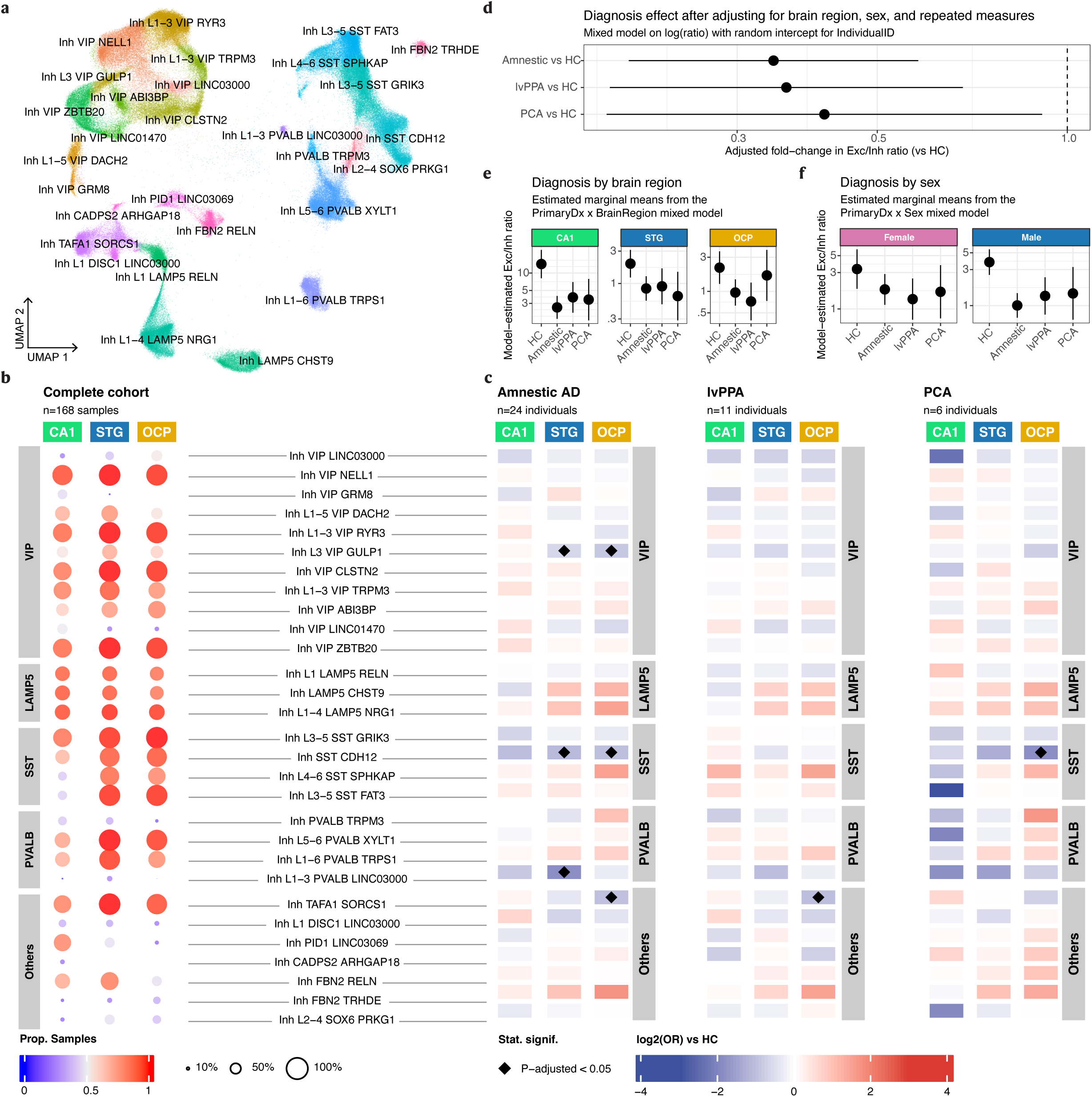
Inhibitory-neuron compositional changes are more limited, but excitatory-to-inhibitory balance is reduced in AD. a, UMAP of 29 inhibitory neuronal subtypes. b, Proportion of region-specific samples in which each inhibitory subtype was detected across the full cohort. c, Relative-abundance modeling of inhibitory subtype changes across diagnoses and regions, visualized as log2-transformed ORs versus HC. Diamonds mark FDR-adjusted P < 0.05. d, Mixed-model estimates of the fold-change in the excitatory-to-inhibitory ratio for each AD syndrome relative to HC. e, Estimated marginal means for the excitatory-to-inhibitory ratio stratified by diagnosis and brain region. f, Estimated marginal means for the excitatory-to-inhibitory ratio stratified by diagnosis and sex.

Relative-abundance modeling showed that inhibitory subtype changes were fewer, more regionally restricted and more mixed in direction than excitatory changes (Fig. 3c). No inhibitory subtype reached significance in CA1. In amnestic AD, STG showed depletion of Inh L1-3 PVALB LINC03000 (OR = 0.27, 95% CI 0.16-0.44, adjusted P = 3.8 × 10^-4^), Inh SST CDH12 (OR = 0.46, 95% CI 0.31-0.68, adjusted P = 0.008) and Inh L3 VIP GULP1 (OR = 0.64, 95% CI 0.49–0.83, adjusted P = 0.022). In OCP, amnestic AD showed depletion of Inh TAFA1 SORCS1 (OR = 0.49, 95% CI 0.35-0.69, adjusted P = 0.003), Inh SST CDH12 (OR = 0.49, 95% CI 0.33-0.71, adjusted P = 0.006) and Inh L3 VIP GULP1 (OR = 0.53, 95% CI 0.36-0.78, adjusted P = 0.015), together with relative enrichment of selected LAMP5-, PVALB- and SST-like states.

In lvPPA, inhibitory depletion was less pronounced and was largely confined to OCP, where Inh TAFA1 SORCS1 remained reduced (OR = 0.48, 95% CI 0.31-0.71, adjusted P = 0.009). In PCA, the clearest inhibitory vulnerability involved OCP Inh SST CDH12 (OR = 0.27, 95% CI 0.12-0.54, adjusted P = 0.011). Thus, although specific inhibitory subtypes were affected in selected regions, the inhibitory compartment did not show the pan-regional depletion pattern observed for Exc NRGN BEX1.

Despite the modest and regionally restricted subtype-level inhibitory changes, the excitatory-to-inhibitory balance was consistently shifted in AD (Fig. 3d). In a linear mixed-effects model, the excitatory-to-inhibitory ratio was lower in amnestic AD (fold-change versus HC = 0.34, 95% CI 0.20-0.58), lvPPA (0.36, 95% CI 0.19-0.68) and PCA (0.41, 95% CI 0.19-0.91). Region-stratified estimated marginal means showed the largest absolute difference in CA1, but the same directional reduction was evident in STG and OCP (Fig. 3e). A similar pattern was observed in both sexes (Fig. 3f), indicating that the reduced excitatory bias of the neuronal compartment was not confined to one sex.

### A conserved synaptic and calcium-handling program distinguishes the vulnerable Exc NRGN BEX1 state

Because Exc NRGN BEX1 showed the most reproducible relative depletion across AD phenotypes and regions, we next asked whether it carried a conserved molecular program that distinguished it from other excitatory subtypes. Restricting analyses to HC nuclei, we performed region-specific pseudobulk differential expression and compared Exc NRGN BEX1 with other regionally prevalent excitatory subtypes in CA1, STG and OCP. Requiring directional concordance across all pairwise contrasts within each region and then intersecting the resulting enriched gene sets across regions identified 72 genes that were commonly enriched in Exc NRGN BEX1 (Fig. 4a).

**Figure 4.**
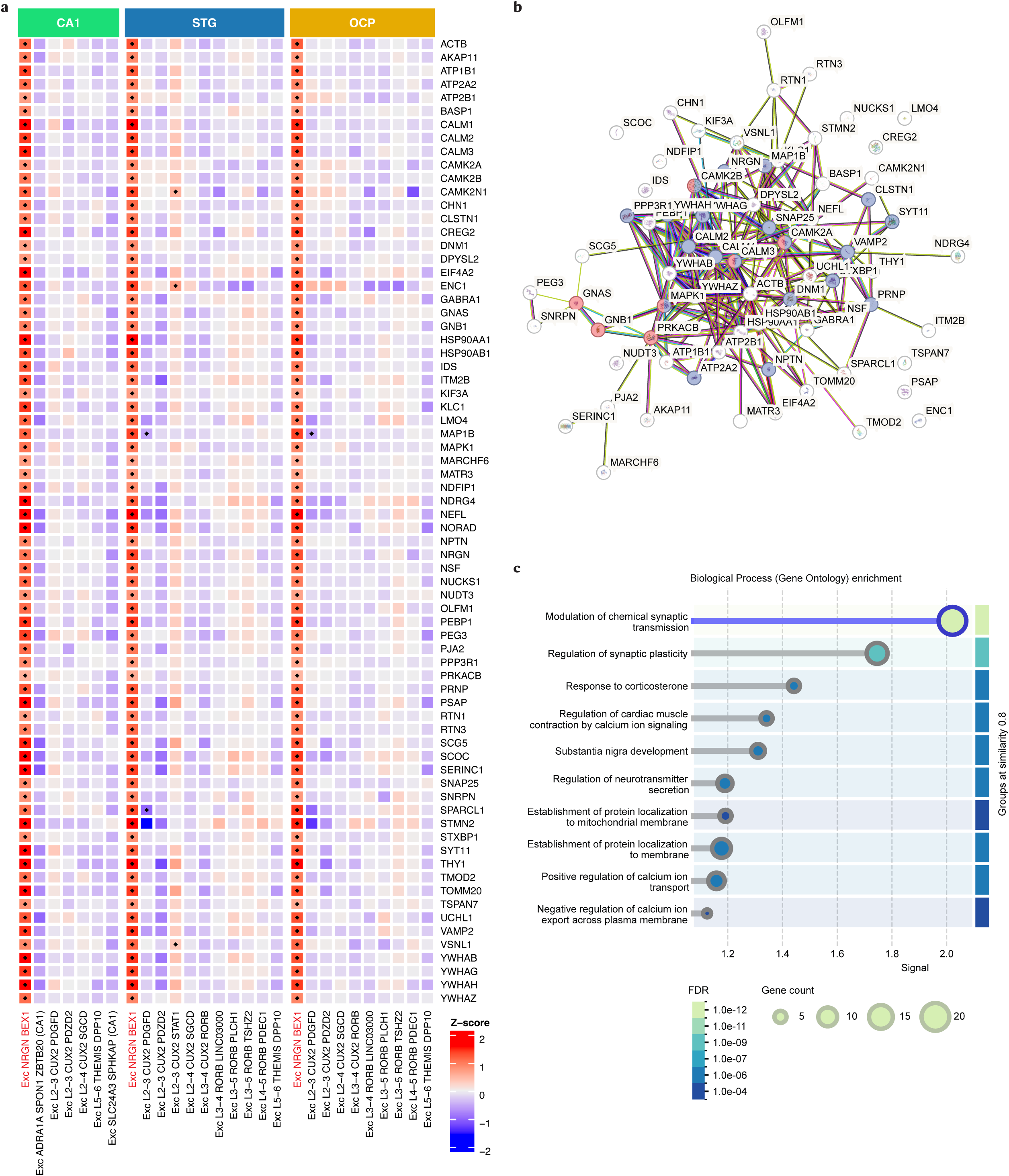
Conserved molecular program of the vulnerable Exc NRGN BEX1 population. a, Heatmap of the 72 genes commonly enriched in Exc NRGN BEX1 across CA1, STG and OCP, displayed across regionally prevalent excitatory subtypes; warmer colors indicate stronger enrichment signal. b, STRING protein–protein interaction network of the shared enriched gene set. c, STRING Gene Ontology biological-process enrichment for the shared BEX1-associated gene set, ranked by FDR and signal strength.

This conserved gene set was enriched for genes involved in synaptic vesicle machinery (for example, SNAP25, VAMP2, STXBP1), calcium and calmodulin signaling (including CAMK2A, CAMK2B, CALM family members), and neuronal structural maintenance (such as MAP1B, NEFL, UCHL1), indicating a coordinated program centered on synaptic function, calcium handling, and neuronal integrity. When visualized across other regionally prevalent vulnerable excitatory subtypes, these genes showed their strongest enrichment in Exc NRGN BEX1 across all three regions (Fig. 4a), indicating that this signature was not broadly shared across all vulnerable excitatory neurons.

STRING analysis of the 72-gene set identified a densely interconnected network (Fig. 4b) and significant enrichment for modulation of chemical synaptic transmission (20 genes, FDR = 1.44 × 10^-12^), regulation of synaptic plasticity (12 genes, FDR = 2.07 × 10^-8^), regulation of neurotransmitter secretion (7 genes, FDR = 3.7 × 10^-5^), establishment of protein localization to membrane (10 genes, FDR = 9.3 × 10^-6^) and positive regulation of calcium ion transport (8 genes, FDR = 3.0 × 10^-5^) (Fig. 4c). Additional enriched terms included response to corticosterone and establishment of protein localization to mitochondrial membrane, further pointing to a transcriptional program shaped by activity-dependent signaling and subcellular trafficking.

### Shared vulnerable neuronal populations are embedded in partially distinct molecular stress programs

Because Exc NRGN BEX1 emerged as a shared vulnerable excitatory population across AD phenotypes, we next asked whether the molecular milieu surrounding this vulnerable state differed by syndrome. We examined curated gene sets related to calcium dysregulation, glial/inflammatory signaling, mitochondrial dysfunction and oxidative stress across the neuronal and glial subpopulations identified above. Calcium-associated changes showed the clearest divergence between phenotypes. Within amnestic AD, this signature was comparatively modest in CA1 but broader in neocortical STG and OCP, indicating broader neocortical transcriptional perturbation rather than a simple recapitulation of the regional NFT gradient (Fig. 5a,b). The same syndrome also showed broader changes in glial/inflammatory genes, including APOE, C1QB, CX3CR1, IL1B, P2RY12 and STAT3, than were apparent in lvPPA or PCA.

**Figure 5.**
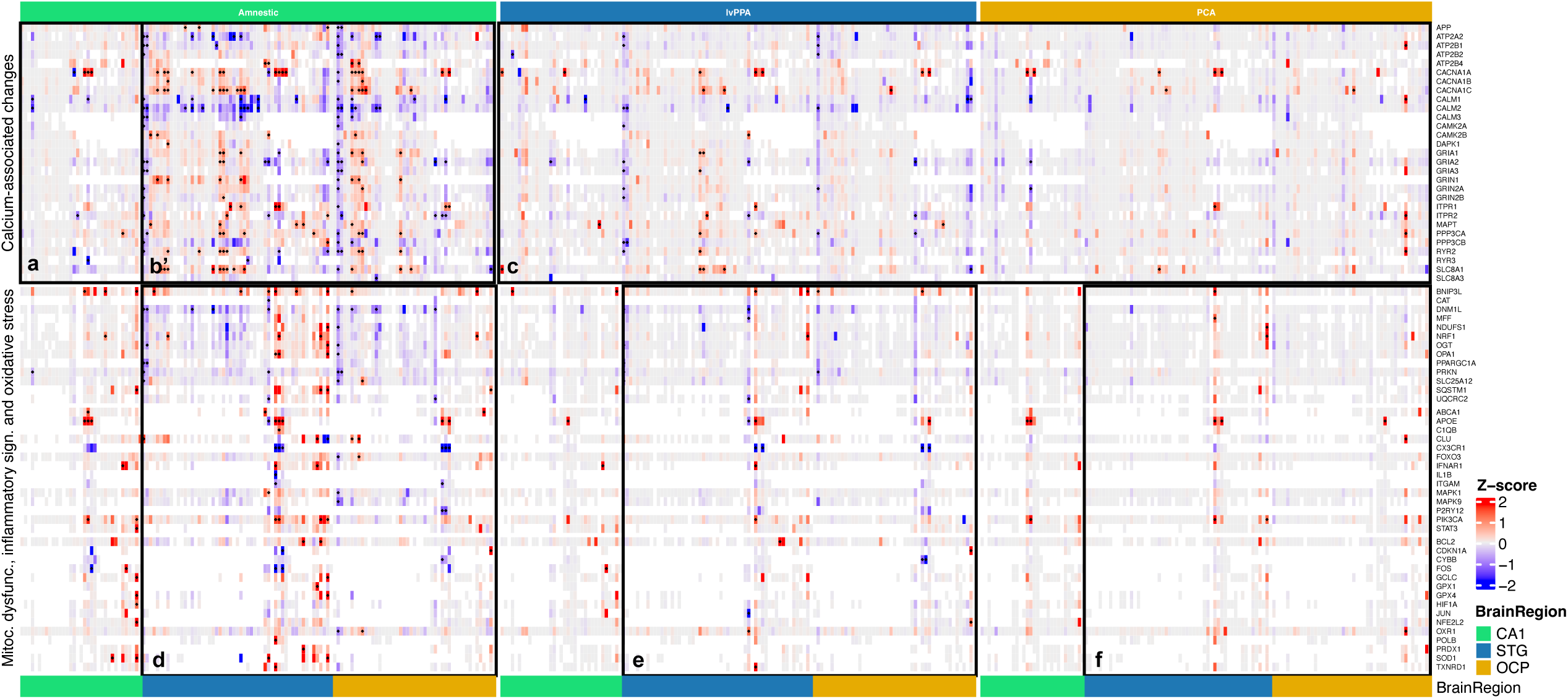
Shared vulnerable neuronal populations are associated with partially distinct molecular stress programs. Heatmaps show standardized differential expression (Z-scores of log2 fold change relative to HC) for curated stress-related gene sets across the neuronal and glial subpopulations defined in Fig. 1. Columns represent snRNA-seq subpopulations grouped by syndrome and annotated by brain region (CA1, STG and OCP); rows represent genes. a and b, within amnestic AD, calcium-associated perturbation is comparatively modest in CA1 but broader in neocortical STG and OCP. c, lvPPA and PCA show sparser calcium-associated signatures than amnestic AD. d, e and f, mitochondrial, neuroinflammatory and oxidative-stress programs are recurrently perturbed across amnestic AD, lvPPA and PCA, consistent with a shared but variably distributed stress axis across phenotypes. Black dots denote gene-subpopulation comparisons reaching adjusted P < 0.05 and absolute log2 fold change > 0.5.

By contrast, lvPPA and PCA showed sparser and more focal calcium-associated signatures despite implicating overlapping vulnerable neuronal populations (Fig. 5c). Mitochondrial, neuroinflammatory and oxidative-stress programs showed greater convergence across phenotypes. Genes involved in mitochondrial dynamics and bioenergetics, such as BNIP3L, DNM1L, OPA1 and PPARGC1A, together with redox-regulatory genes including GPX4, NFE2L2, PRDX1 and SOD1, were recurrently perturbed in amnestic AD, lvPPA and PCA, although their magnitude and cellular distribution varied across subpopulations and regions (Fig. 5d-f). Taken together, these data suggest that the same vulnerable neuronal populations are embedded in partially distinct syndrome-associated molecular contexts, with stronger calcium-associated transcriptional perturbation in amnestic AD and more variably distributed metabolic, oxidative and glial-inflammatory changes across all phenotypes.

## Discussion

By leveraging neuropathologically matched typical and atypical AD phenotypes, this study identifies a conserved excitatory neuronal state that is selectively vulnerable across clinically distinct forms of AD. The central finding is the reproducible depletion of Exc NRGN BEX1: it was significantly reduced across CA1, STG and OCP in amnestic AD and across STG and OCP in lvPPA, with parallel histological validation in STG. This state aligns closely with previously described AD-vulnerable excitatory populations. In our cross-dataset correlations, Exc NRGN BEX1 showed its strongest correspondence to the vulnerable Exc.s2 states reported by Leng and colleagues in the entorhinal cortex (ρ = 0.803) (Supplementary Fig. 4) [8]. It also showed strong similarity to the NRGN excitatory populations in the multiregional atlas from Mathys and colleagues, with the highest correlation observed for prefrontal-cortex Exc NRGN (ρ = 0.874) and similarly high correlations across angular gyrus, midtemporal cortex, entorhinal cortex and hippocampus (ρ = 0.863-0.872) (Supplementary Fig. 5) [10]. Together with prior work showing preferential excitatory-neuron susceptibility to tau pathology and network-level molecular determinants of selective vulnerability, these convergences argue that the BEX1-positive NRGN population identified here represents a conserved AD-vulnerable excitatory program rather than a cohort-specific state [20, 21].

We next asked why this population might be vulnerable. The conserved BEX1-associated signature was dominated by genes involved in synaptic vesicle release, membrane excitability, calcium/calmodulin signaling and structural plasticity, and the two most significant biological-process enrichments were modulation of chemical synaptic transmission and regulation of synaptic plasticity. A parsimonious interpretation is that neurons specialized for high activity-dependent plasticity may be especially sensitive to tau-associated dysfunction. Neurogranin is a calmodulin-binding postsynaptic protein that modulates synaptic plasticity through calcium-dependent signaling, and recent work in vulnerable neurons has linked cell-autonomous regulation of neuronal excitability to tau accumulation and microglia-mediated degeneration [22,23]. In this framework, the same molecular architecture that supports synaptic gain may also impose a higher burden on calcium homeostasis, vesicle dynamics and proteostatic control, thereby narrowing the margin between physiological specialization and pathological collapse.

This mechanistic interpretation remains provisional, however, because our cross-sectional design does not distinguish pre-existing neuronal properties from adaptive or end-stage transcriptional responses.

Inhibitory neurons showed a more limited and regionally restricted pattern of compositional change. Specific interneuron states were depleted in STG and OCP, but inhibitory subtype changes were mixed in direction and did not reproduce the cross-region consistency seen for Exc NRGN BEX1. Nevertheless, the excitatory-to-inhibitory ratio was reduced across all AD phenotypes, indicating that preferential depletion of excitatory neurons is sufficient to shift the balance of the neuronal compartment even when interneuron loss is modest. Because this ratio is derived from class proportions in snRNA-seq data, it should be interpreted as a compositional index rather than as a direct physiological readout of circuit excitation-inhibition. Even so, the result is consistent with patient-based evidence that excitatory and inhibitory parameters relate differently to tau and amyloid burden, supporting the idea that these neuronal compartments are not affected symmetrically in AD [24].

Although amnestic AD, lvPPA and PCA converge on depletion of the same vulnerable excitatory population, the surrounding transcriptional programs are not equivalent. In our curated pathway analysis, amnestic AD showed broader calcium-associated perturbation, particularly in neocortical regions, whereas mitochondrial, glial/inflammatory and oxidative-stress signatures recurred across all phenotypes. Because these heatmaps derive from targeted gene sets in a cross-sectional dataset and do not directly compare syndromes with one another, we interpret them as hypothesis-generating evidence for partially distinct molecular contexts acting on a shared vulnerable neuronal target rather than as definitive proof of syndrome-specific upstream mechanisms. This point is especially important because, within amnestic AD, the calcium-associated signature was comparatively modest in CA1 and broader in STG and OCP, indicating that these transcriptional perturbations do not simply recapitulate the regional NFT hierarchy. In addition, the broader detectable signal in amnestic AD should be interpreted cautiously, because unequal group sizes may have increased power to detect transcriptomic changes in that group relative to lvPPA and PCA. Taken together, these findings are more consistent with partially distinct molecular environments converging on a shared vulnerable neuronal substrate than with entirely different neuronal targets across syndromes.

Several limitations should be considered. First, the relative-abundance models are compositional and therefore detect depletion or enrichment within a parent class rather than absolute neuronal counts. Second, the study is cross-sectional and therefore cannot determine whether observed transcriptional differences represent premorbid susceptibility, active adaptation, or selective survival of particular cell states. Third, the PCA cohort was comparatively small, which probably limited power to detect subtler excitatory depletion in that syndrome. Fourth, orthogonal histological validation was restricted to STG. Fifth, all cases were APOE ε3/ε3 carriers, which reduced genetic heterogeneity but also narrows generalizability, especially because APOE status influences medial temporal atrophy and tau deposition in atypical AD [25]. Despite these limitations, the present study connects regionally patterned NFT burden to cell-intrinsic vulnerability programs across clinically distinct AD phenotypes. A conserved NRGN-BEX1 excitatory state emerges as a cross-phenotype locus of vulnerability and a tractable entry point for mechanistic studies aimed at explaining why tau-rich cortical and hippocampal networks show selective vulnerability and relative depletion of specific neuronal states in AD.

## Acknowledgments

This work was supported by grants from Tau Consortium/Rainwater Charity Foundation, NIA R01AG060477, NIA R01 AG064314, NIA K24 AG053435 (Grinberg).

## Supplementary materials

**Supplementary Figure 1 | Cross-region and neuropathological group correlations.**

Heatmap showing Spearman correlations of major cell classes (excitatory neurons, inhibitory neurons, glial cells, and vascular cells) across brain regions and neuropathological groups.

**Supplementary Figure 2 | Sub-neuronal population module scores.**

Scatter plots of module scores across sub-neuronal populations, together with fitted regression models (see Methods).

**Supplementary Figures 3-5 | Cross-dataset correlation of cell type subpopulations.**

Heatmaps showing Spearman correlations of cell type subpopulations across independent datasets. Supplementary Figure 3 corresponds to the multi–brain region Allen Brain Institute dataset; Supplementary Figure 4 to the entorhinal cortex and superior temporal gyrus dataset from Leng et al. (2021); and Supplementary Figure 5 to the multiregional dataset from Mathys et al. (2024).

