## Supplementary Table 1 for "Single-nucleus transcriptomics identifies a shared vulnerable excitatory neuronal population across typical and atypical Alzheimer’s disease"

Supplementary Table 1. **Detailed cohort characteristics and neuropathological features**

| Case | Sex | AAD | Neuro-pathologic Group | Braak Stage | ABC | APOE | PMI | CA1 | STG | OCP |
|---|---|---|---|---|---|---|---|---|---|---|
| Case #1 | Male | 55.5 | HC | 0 | A≤1B≤1C≤1 | 3/3 | 18.2 | x | x | |
| Case #2 | Male | 67 | HC | 0 | A≤1B≤1C≤1 | 3/3 | 14.2 | x | x | |
| Case #3 | Male | 59.9 | HC | 0 | A≤1B≤1C≤1 | 3/3 | 15.1 | | | x |
| Case #4 | Male | 68.4 | HC | 0 | A≤1B≤1C≤1 | 3/3 | 14.1 | | | x |
| Case #5 | Male | 53.8 | HC | 0 | A≤1B≤1C≤1 | 3/3 | 14 | | | x |
| Case #6 | Female | 82 | HC | 0 | A≤1B≤1C≤1 | 3/3 | 36 | x | x | x |
| Case #7 | Male | 69 | HC | 0 | A≤1B≤1C≤1 | 3/3 | NA | x | x | x |
| Case #8 | Female | 80 | HC | 0 | A≤1B≤1C≤1 | 3/3 | NA | x | x | x |
| Case #9 | Male | NA | HC | 0 | A≤1B≤1C≤1 | 3/3 | 19.4 | x | x | x |
| Case #10 | Male | 55.5 | HC | I | A≤1B≤1C≤1 | 3/3 | 20.1 | | x | |
| Case #11 | Male | 61.7 | HC | I | A≤1B≤1C≤1 | 3/3 | 8.2 | | | x |
| Case #12 | Male | 65.9 | HC | I | A≤1B≤1C≤1 | 3/3 | 11.2 | | x | |
| Case #13 | Female | 50.6 | HC | I | A≤1B≤1C≤1 | 3/3 | 15.6 | x | x | x |
| Case #14 | Female | 59.1 | HC | I | A≤1B≤1C≤1 | 3/3 | 13.4 | | | x |
| Case #15 | Male | 71.2 | HC | I | A≤1B≤1C≤1 | 3/3 | 11.9 | | x | |
| Case #16 | Male | 61.9 | HC | I | A≤1B≤1C≤1 | 3/3 | 13.7 | x | x | |
| Case #17 | Female | 64.8 | HC | I | A≤1B≤1C≤1 | 3/3 | 11.2 | x | x | |
| Case #18 | Male | 59.3 | HC | I | A≤1B≤1C≤1 | 3/3 | 12.2 | x | x | |
| Case #19 | Male | 63.7 | HC | I | A≤1B≤1C≤1 | 3/3 | 11.9 | x | x | |
| Case #20 | Female | 54.4 | HC | I | A≤1B≤1C≤1 | 3/3 | NA | | x | x |
| Case #21 | Female | 85.3 | HC | I | A≤1B≤1C≤1 | 3/3 | 6 | | x | x |
| Case #22 | Male | 67.1 | HC | II | A≤1B≤1C≤1 | 3/3 | 9.7 | x | x | |
| Case #23 | Male | 50.7 | HC | II | A≤1B≤1C≤1 | 3/3 | 14.5 | | | x |
| Case #24 | Male | 65.9 | HC | II | A≤1B≤1C≤1 | 3/3 | 12.3 | | | x |
| Case #25 | Male | 57.7 | HC | II | A≤1B≤1C≤1 | 3/3 | 19 | x | x | |
| Case #26 | Male | 78.1 | HC | II | A≤1B≤1C≤1 | 3/3 | 8 | x | x | x |
| Case #27 | Male | 82 | HC | II | A≤1B≤1C≤1 | 3/3 | NA | | | x |
| Case #28 | Male | 57 | Amnestic | V | A3B3C3 | 3/3 | 6 | x | x | x |
| Case #29 | Male | 67.4 | lvPPA | V | A3B3C3 | 3/3 | NA | x | x | x |
| Case #30 | Male | 79 | Amnestic | VI | A3B3C3 | 3/3 | 11 | x | x | x |
| Case #31 | Female | 64 | lvPPA | VI | A3B3C3 | 3/3 | 3 | x | x | x |
| Case #32 | Female | 69 | lvPPA | VI | A3B3C3 | 3/3 | 2.6 | x | x | x |
| Case #33 | Female | 75 | Amnestic | VI | A3B3C3 | 3/3 | NA | x | x | x |
| Case #34 | Male | 63 | Amnestic | VI | A3B3C3 | 3/3 | 4.5 | x | x | x |
| Case #35 | Female | 67 | Amnestic | VI | A3B3C3 | 3/3 | NA | x | x | x |
| Case #36 | Female | 70 | Amnestic | VI | A3B3C3 | 3/3 | 8.2 | x | x | x |
| Case #37 | Male | 75 | Amnestic | VI | A3B3C3 | 3/3 | NA | x | x | x |
| Case #38 | Male | 54.6 | Amnestic | VI | A3B3C3 | 3/3 | 17 | x | x | x |
| Case #39 | Male | 65 | Amnestic | VI | A3B3C3 | 3/3 | 23 | x | x | x |
| Case #40 | Male | 73.3 | lvPPA | VI | A3B3C3 | 3/3 | 7 | x | x | x |
| Case #41 | Male | 67.6 | PCA | VI | A3B3C3 | 3/3 | NA | x | x | x |
| Case #42 | Female | 58.9 | Amnestic | VI | A3B3C3 | 3/3 | NA | | x | x |
| Case #43 | Female | 71.7 | Amnestic | VI | A3B3C3 | 3/3 | 9 | | x | x |
| Case #44 | Female | 65.3 | Amnestic | VI | A3B3C3 | 3/3 | 7 | | x | x |
| Case #45 | Female | 72.3 | PCA | VI | A3B3C3 | 3/3 | 11 | x | x | x |
| Case #46 | Male | 62.2 | lvPPA | VI | A3B3C3 | 3/3 | 5 | x | x | x |
| Case #47 | Male | 61.2 | PCA | VI | A3B3C3 | 3/3 | 4 | x | x | x |
| Case #48 | Male | 66.4 | Amnestic | VI | A3B3C3 | 3/3 | 10 | x | x | x |
| Case #49 | Female | 68.6 | Amnestic | VI | A3B3C3 | 3/3 | 3 | x | x | x |

| | | | | | | | | | | |
|---|---|---|---|---|---|---|---|---|---|---|
| Case #50 | Male | 71 | Amnestic | VI | A3B3C3 | 3/3 | NA | | x | x |
| Case #51 | Male | 66.6 | lvPPA | VI | A3B3C3 | 3/3 | NA | x | x | x |
| Case #52 | Male | 64 | Amnestic | VI | A3B3C3 | 3/3 | 6 | x | x | x |
| Case #53 | Male | 70 | Amnestic | VI | A3B3C3 | 3/3 | NA | x | x | x |
| Case #54 | Male | 63 | lvPPA | VI | A3B3C3 | 3/3 | 17.8 | x | x | x |
| Case #55 | Female | 69 | Amnestic | VI | A3B3C3 | 3/3 | 12.9 | x | x | x |
| Case #56 | Male | 62 | lvPPA | VI | A3B3C3 | 3/3 | 9.9 | x | x | x |
| Case #57 | Male | 66 | Amnestic | VI | A3B3C3 | 3/3 | 25 | x | x | x |
| Case #58 | Female | 66 | lvPPA | VI | A3B3C3 | 3/3 | 26.7 | x | x | x |
| Case #59 | Male | 68 | Amnestic | VI | A3B3C3 | 3/3 | 9.3 | x | x | x |
| Case #60 | Male | 64 | Amnestic | VI | A3B3C3 | 3/3 | 6.8 | x | x | x |
| Case #61 | Male | 62 | Amnestic | VI | A3B3C3 | 3/3 | 6 | x | x | x |
| Case #62 | Female | 64 | Amnestic | VI | A3B3C3 | 3/3 | 8.9 | x | x | x |
| Case #63 | Female | 68 | Amnestic | VI | A3B3C3 | 3/3 | 18 | x | x | x |
| Case #64 | Female | 64 | lvPPA | VI | A3B3C3 | 3/3 | 20.8 | x | x | x |
| Case #65 | Male | 71 | PCA | VI | A3B3C3 | 3/3 | 6.5 | x | x | x |
| Case #66 | Female | 72 | PCA | VI | A3B3C3 | 3/3 | 7.4 | x | x | x |
| Case #67 | Female | 52 | lvPPA | VI | A3B3C3 | 3/3 | 52 | x | x | x |
| Case #68 | Female | 80 | PCA | VI | A3B3C3 | 3/3 | 19.9 | x | x | x |

AAD = Age at Death; ABC = ABC dementia scale classes for AD patients; APOE = Apolipoprotein E; PMI = post-mortem interval; CA1 = Cornu Ammonis sector 1 of Hippocampus. STG = superior temporal gyrus; OCP = occipital cortex; HC = healthy control; lvPPA = logopenic variant Primary Progressive Aphasia; PCA = Posterior Cortical Atrophy.
