## Supplementary Figures for "Single-nucleus transcriptomics identifies a shared vulnerable excitatory neuronal population across typical and atypical Alzheimer’s disease"

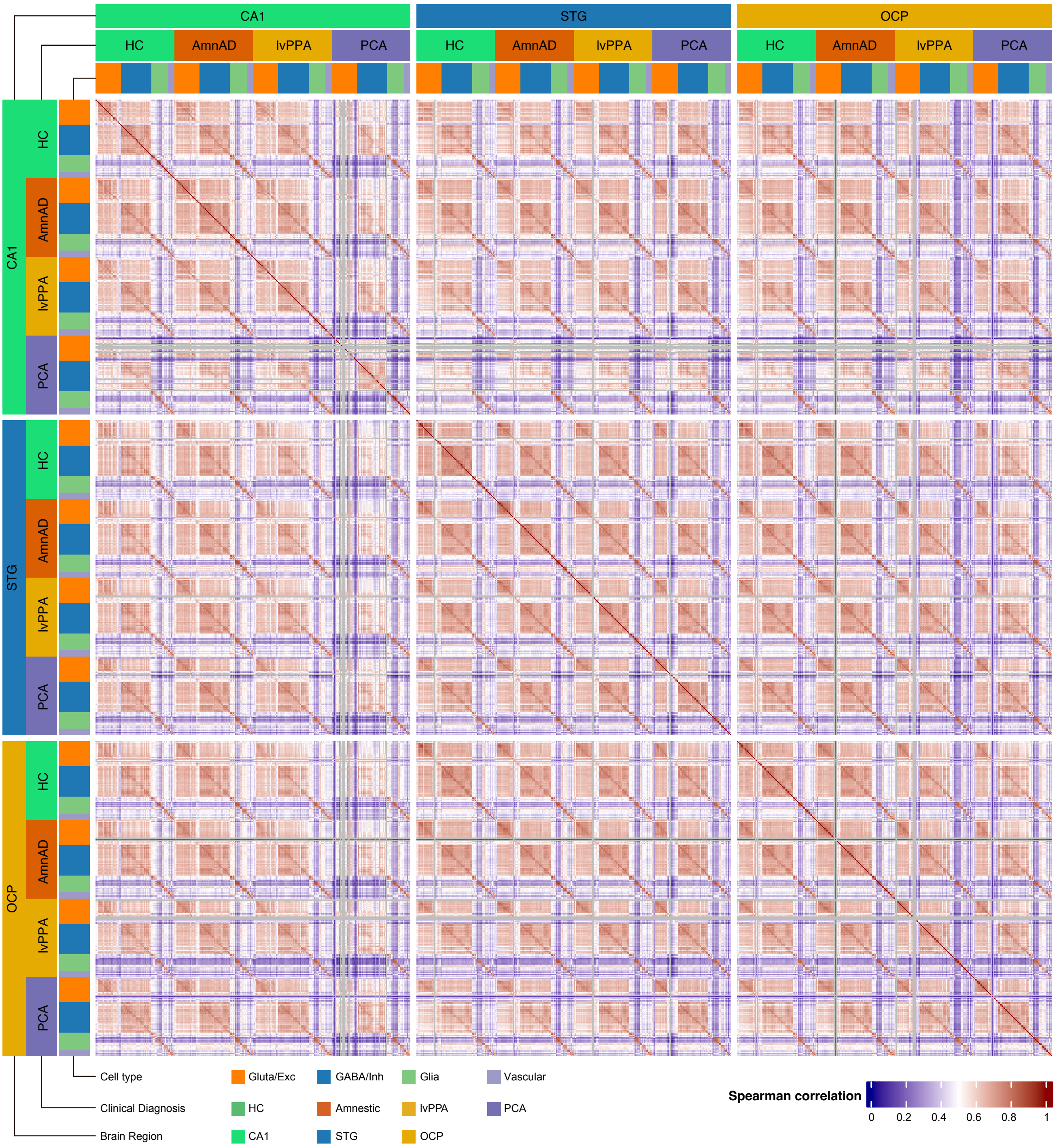
CA1
STG
OCP
HC
AmnAD
lvPPA
PCA
Cell type
Gluta/Exc
GABA/Inh
Glia
Vascular
Clinical Diagnosis
HC
Amnestic
lvPPA
PCA
Brain Region
CA1
STG
OCP
Spearman correlation
0
0.2
0.4
0.6
0.8
1

### Supplementary Figure 1 | Cross-region and neuropathological group correlations.

Heatmap showing Spearman correlations of major cell classes (excitatory neurons, inhibitory neurons, glial cells, and vascular cells) across brain regions and neuropathological groups.

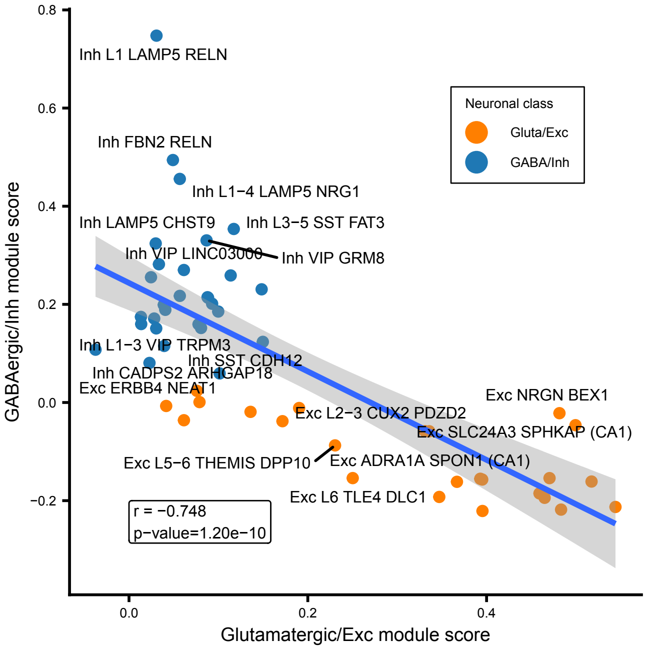

Inh L1 LAMP5 RELN
Inh FBN2 RELN
Inh L1−4 LAMP5 NRG1
Inh LAMP5 CHST9
Inh L3−5 SST FAT3
Inh VIP LINC03000
Inh VIP GRM8
Inh L1−3 VIP TRPM3
Inh SST CDH12
Inh CADPS2 ARHGAP18
Exc ERBB4 NEAT1
Exc NRGN BEX1
Exc L2−3 CUX2 PDZD2
Exc SLC24A3 SPHKAP (CA1)
Exc L5−6 THEMIS DPP10
Exc ADRA1A SPON1 (CA1)
Exc L6 TLE4 DLC1
Neuronal class
Gluta/Exc
GABA/Inh
r = −0.748
p−value=1.20e−10
0.8
0.6
0.4
0.2
0.0
−0.2
0.0
0.2
0.4
GABAergic/Inh module score
Glutamatergic/Exc module score

### Supplementary Figure 2 | Sub-neuronal population module scores.

Scatter plots of module scores across sub-neuronal populations, together with fitted regression models (see Methods).

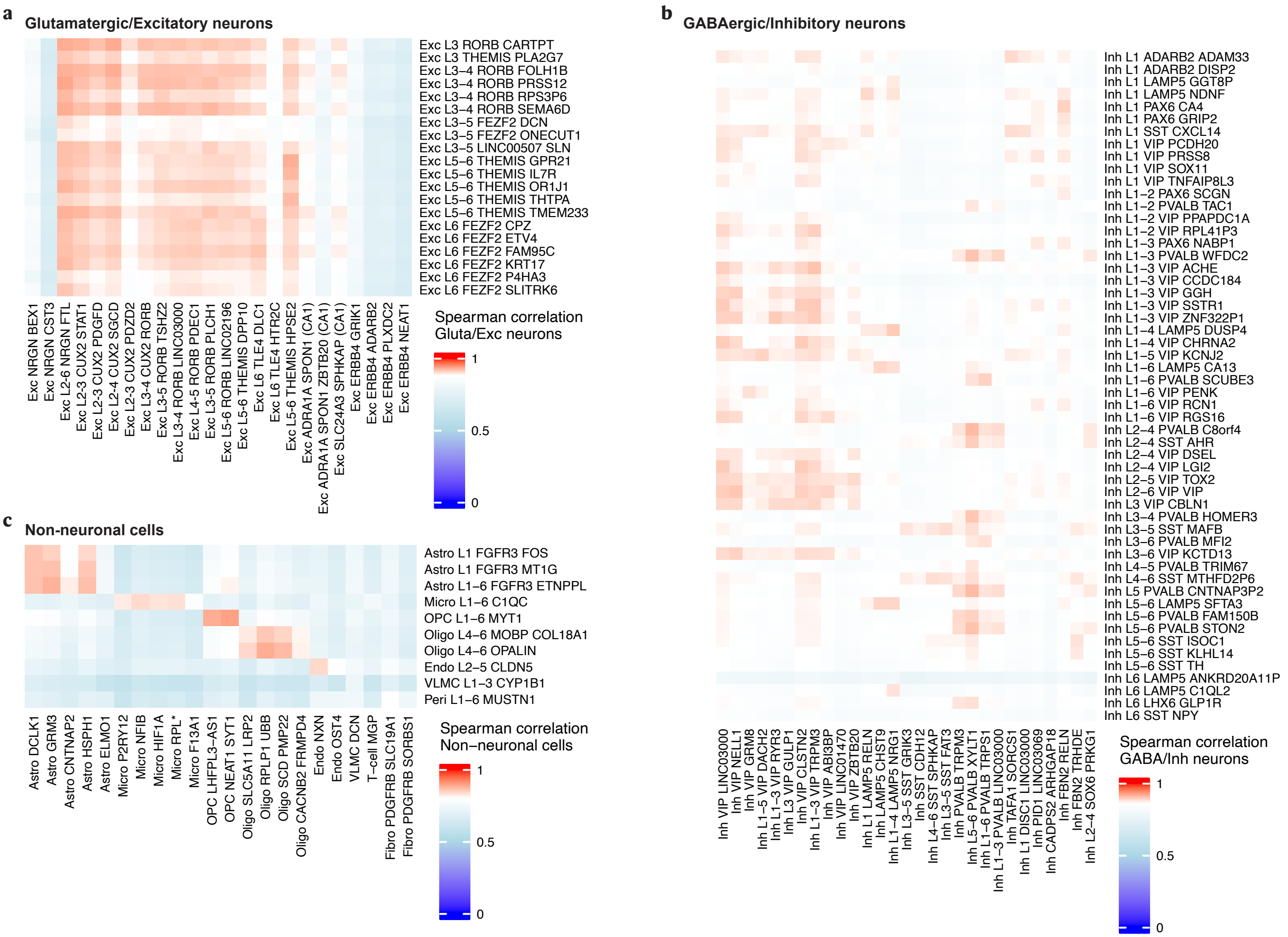
a
Glutamatergic/Excitatory neurons
Exc L3 RORB CARTPT
Exc L3 THEMIS PLA2G7
Exc L3–4 RORB FOLH1B
Exc L3–4 RORB PRSS12
Exc L3–4 RORB RPS3P6
Exc L3–4 RORB SEMA6D
Exc L3–5 FEZF2 DCN
Exc L3–5 FEZF2 ONECUT1
Exc L3–5 LINC00507 SLN
Exc L5–6 THEMIS GPR21
Exc L5–6 THEMIS IL7R
Exc L5–6 THEMIS OR1J1
Exc L5–6 THEMIS THTPA
Exc L5–6 THEMIS TMEM233
Exc L6 FEZF2 CPZ
Exc L6 FEZF2 ETV4
Exc L6 FEZF2 FAM95C
Exc L6 FEZF2 KRT17
Exc L6 FEZF2 P4HA3
Exc L6 FEZF2 SLITRK6
Exc NRGN BEX1
Exc NRGN CST3
Exc L2-6 NRGN FTL
Exc L2-3 CUX2 STAT1
Exc L2-3 CUX2 PDGFD
Exc L2-4 CUX2 SGCD
Exc L2-3 CUX2 PDZD2
Exc L3-4 CUX2 RORB
Exc L3-5 RORB TSHZ2
Exc L3-4 RORB LINC03000
Exc L4-5 RORB PDEC1
Exc L3-5 RORB PLCH1
Exc L5-6 RORB LINC02196
Exc L5-6 THEMIS DPP10
Exc L6 TLE4 DLC1
Exc L6 TLE4 HTR2C
Exc L5-6 THEMIS HPSE2
Exc ADRA1A SPON1 (CA1)
Exc ADRA1A SPON1 ZBTB20 (CA1)
Exc SLC24A3 SPHKAP (CA1)
Exc ERBB4 GRIK1
Exc ERBB4 ADARB2
Exc ERBB4 PLXDC2
Exc ERBB4 NEAT1
Spearman correlation
Gluta/Exc neurons
1
0.5
0
b
GABAergic/Inhibitory neurons
Inh L1 ADARB2 ADAM33
Inh L1 ADARB2 DISP2
Inh L1 LAMP5 GGT8P
Inh L1 LAMP5 NDNF
Inh L1 PAX6 CA4
Inh L1 PAX6 GRIP2
Inh L1 SST CXCL14
Inh L1 VIP PCDH20
Inh L1 VIP PRSS8
Inh L1 VIP SOX11
Inh L1 VIP TNFAIP8L3
Inh L1–2 PAX6 SCGN
Inh L1–2 PVALB TAC1
Inh L1–2 VIP PPAPDC1A
Inh L1–2 VIP RPL41P3
Inh L1–3 PAX6 NABP1
Inh L1–3 PVALB WFDC2
Inh L1–3 VIP ACHE
Inh L1–3 VIP CCDC184
Inh L1–3 VIP GGH
Inh L1–3 VIP SSTR1
Inh L1–3 VIP ZNF322P1
Inh L1–4 LAMP5 DUSP4
Inh L1–4 VIP CHRNA2
Inh L1–5 VIP KCNJ2
Inh L1–6 LAMP5 CA13
Inh L1–6 PVALB SCUBE3
Inh L1–6 VIP PENK
Inh L1–6 VIP RCN1
Inh L1–6 VIP RGS16
Inh L2–4 PVALB C8orf4
Inh L2–4 SST AHR
Inh L2–4 VIP DSEL
Inh L2–4 VIP LGI2
Inh L2–5 VIP TOX2
Inh L2–6 VIP VIP
Inh L3 VIP CBLN1
Inh L3–4 PVALB HOMER3
Inh L3–5 SST MAFB
Inh L3–6 PVALB MFI2
Inh L3–6 VIP KCTD13
Inh L4–5 PVALB TRIM67
Inh L4–6 SST MTHFD2P6
Inh L5 PVALB CNTNAP3P2
Inh L5–6 LAMP5 SFTA3
Inh L5–6 PVALB FAM150B
Inh L5–6 PVALB STON2
Inh L5–6 SST ISOC1
Inh L5–6 SST KLHL14
Inh L5–6 SST TH
Inh L6 LAMP5 ANKRD20A11P
Inh L6 LAMP5 C1QL2
Inh L6 LHX6 GLP1R
Inh L6 SST NPY
Inh VIP LINC03000
Inh VIP NELL1
Inh VIP GRM8
Inh L1–5 VIP DACH2
Inh L1–3 VIP RYR3
Inh L3 VIP GULP1
Inh VIP CLSTN2
Inh L1–3 VIP TRPM3
Inh VIP ABI3BP
Inh VIP LINC01470
Inh VIP ZBTB20
Inh L1 LAMP5 RELN
Inh LAMP5 CHST9
Inh L1–4 LAMP5 NRG1
Inh L3–5 SST GRIK3
Inh SST CDH12
Inh L4–6 SST SPHKAP
Inh L3–5 SST FAT3
Inh PVALB TRPM3
Inh L5–6 PVALB XYLT1
Inh L1–6 PVALB TRPS1
Inh L1–3 PVALB LINC03000
Inh TAFA1 SORCS1
Inh L1 DISC1 LINC03000
Inh PID1 LINC03069
Inh CADPS2 ARHGAP18
Inh FBN2 RELN
Inh FBN2 TRHDE
Inh L2–4 SOX6 PRKG1
Spearman correlation
GABA/Inh neurons
1
0.5
0
c
Non-neuronal cells
Astro L1 FGFR3 FOS
Astro L1 FGFR3 MT1G
Astro L1–6 FGFR3 ETNPPL
Micro L1–6 C1QC
OPC L1–6 MYT1
Oligo L4–6 MOBP COL18A1
Oligo L4–6 OPALIN
Endo L2–5 CLDN5
VLMC L1–3 CYP1B1
Peri L1–6 MUSTN1
Astro DCLK1
Astro GRM3
Astro CNTNAP2
Astro HSPH1
Astro ELMO1
Micro P2RY12
Micro NFIB
Micro HIF1A
Micro RPL*
Micro F13A1
OPC LHFPL3-AS1
OPC NEAT1 SYT1
Oligo SLC5A11 LRP2
Oligo RPLP1 UBB
Oligo SCD PMP22
Oligo CACNB2 FRMPD4
Endo NXN
Endo OST4
VLMC DCN
T-cell MGP
Fibro PDGFRB SLC19A1
Fibro PDGFRB SORBS1
Spearman correlation
Non–neuronal cells
1
0.5
0

### Supplementary Figures 3-5 | Cross-dataset correlation of cell type subpopulations.

Heatmaps showing Spearman correlations of cell type subpopulations across independent datasets. Supplementary Figure 3 corresponds to the multi–brain region Allen Brain Institute dataset; Supplementary Figure 4 to the entorhinal cortex and superior temporal gyrus dataset from Leng et al. (2021); and Supplementary Figure 5 to the multiregional dataset from Mathys et al. (2024).

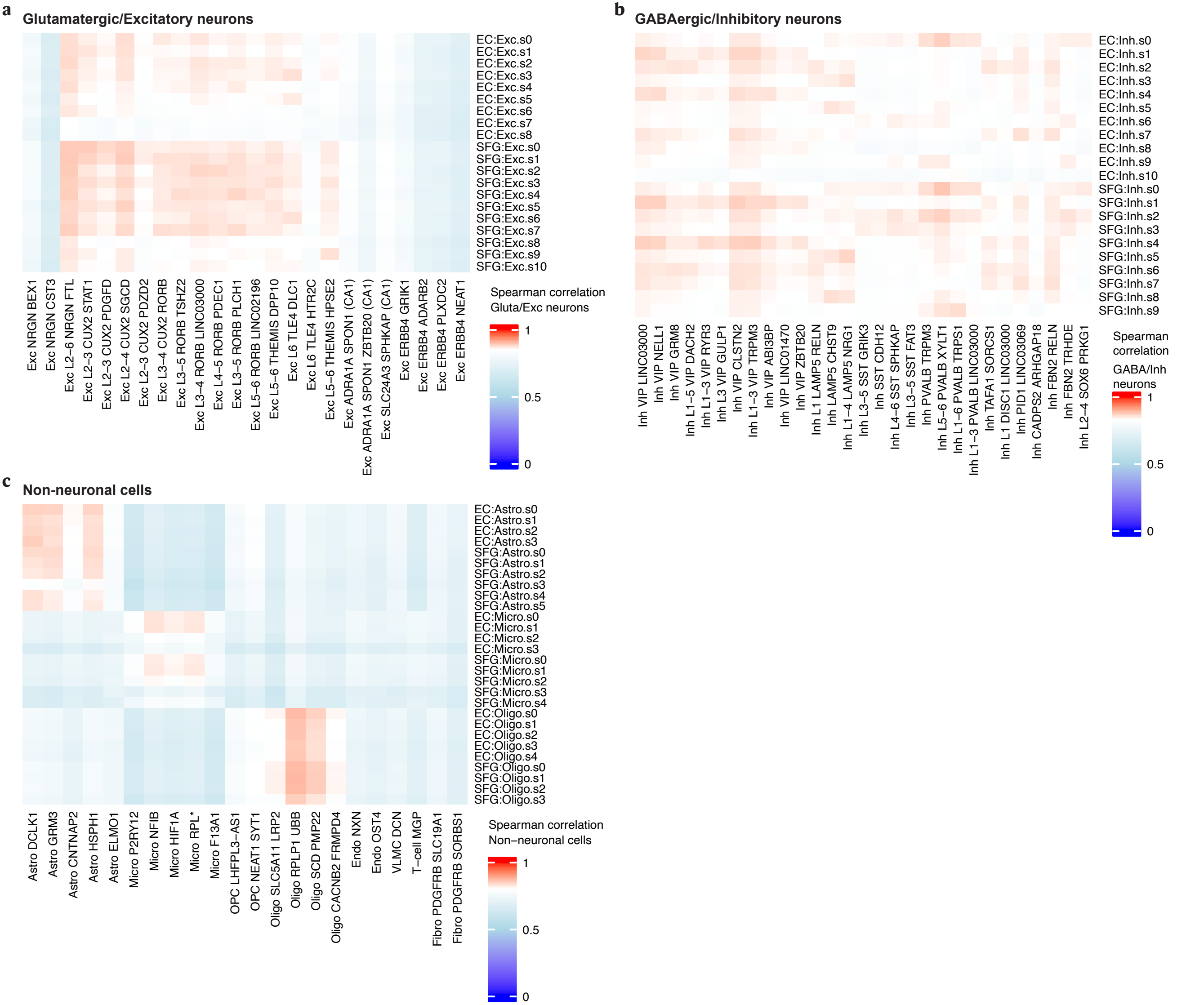
a
Glutamatergic/Excitatory neurons
EC:Exc.s0
EC:Exc.s1
EC:Exc.s2
EC:Exc.s3
EC:Exc.s4
EC:Exc.s5
EC:Exc.s6
EC:Exc.s7
EC:Exc.s8
SFG:Exc.s0
SFG:Exc.s1
SFG:Exc.s2
SFG:Exc.s3
SFG:Exc.s4
SFG:Exc.s5
SFG:Exc.s6
SFG:Exc.s7
SFG:Exc.s8
SFG:Exc.s9
SFG:Exc.s10
Exc NRGN BEX1
Exc NRGN CST3
Exc L2−6 NRGN FTL
Exc L2−3 CUX2 STAT1
Exc L2−3 CUX2 PDGFD
Exc L2−4 CUX2 SGCD
Exc L2−3 CUX2 PDZD2
Exc L3−4 CUX2 RORB
Exc L3−5 RORB TSHZ2
Exc L3−4 RORB LINC03000
Exc L4−5 RORB PDEC1
Exc L3−5 RORB PLCH1
Exc L5−6 RORB LINC02196
Exc L5−6 THEMIS DPP10
Exc L6 TLE4 DLC1
Exc L6 TLE4 HTR2C
Exc L5−6 THEMIS HPSE2
Exc ADRA1A SPON1 (CA1)
Exc ADRA1A SPON1 ZBTB20 (CA1)
Exc SLC24A3 SPHKAP (CA1)
Exc ERBB4 GRIK1
Exc ERBB4 ADARB2
Exc ERBB4 PLXDC2
Exc ERBB4 NEAT1
Spearman correlation
Gluta/Exc neurons
1
0.5
0
b
GABAergic/Inhibitory neurons
EC:Inh.s0
EC:Inh.s1
EC:Inh.s2
EC:Inh.s3
EC:Inh.s4
EC:Inh.s5
EC:Inh.s6
EC:Inh.s7
EC:Inh.s8
EC:Inh.s9
EC:Inh.s10
SFG:Inh.s0
SFG:Inh.s1
SFG:Inh.s2
SFG:Inh.s3
SFG:Inh.s4
SFG:Inh.s5
SFG:Inh.s6
SFG:Inh.s7
SFG:Inh.s8
SFG:Inh.s9
Inh VIP LINC03000
Inh VIP NELL1
Inh VIP GRM8
Inh L1−5 VIP DACH2
Inh L1−3 VIP RYR3
Inh L3 VIP GULP1
Inh VIP CLSTN2
Inh L1−3 VIP TRPM3
Inh VIP ABI3BP
Inh VIP LINC01470
Inh VIP ZBTB20
Inh L1 LAMP5 RELN
Inh LAMP5 CHST9
Inh L1−4 LAMP5 NRG1
Inh L3−5 SST GRIK3
Inh SST CDH12
Inh L4−6 SST SPHKAP
Inh L3−5 SST FAT3
Inh PVALB TRPM3
Inh L5−6 PVALB XYLT1
Inh L1−6 PVALB TRPS1
Inh L1−3 PVALB LINC03000
Inh TAFA1 SORCS1
Inh L1 DISC1 LINC03000
Inh PID1 LINC03069
Inh CADPS2 ARHGAP18
Inh FBN2 RELN
Inh FBN2 TRHDE
Inh L2−4 SOX6 PRKG1
Spearman
correlation
GABA/Inh
neurons
1
0.5
0
c
Non-neuronal cells
EC:Astro.s0
EC:Astro.s1
EC:Astro.s2
EC:Astro.s3
SFG:Astro.s0
SFG:Astro.s1
SFG:Astro.s2
SFG:Astro.s3
SFG:Astro.s4
SFG:Astro.s5
EC:Micro.s0
EC:Micro.s1
EC:Micro.s2
EC:Micro.s3
SFG:Micro.s0
SFG:Micro.s1
SFG:Micro.s2
SFG:Micro.s3
SFG:Micro.s4
EC:Oligo.s0
EC:Oligo.s1
EC:Oligo.s2
EC:Oligo.s3
EC:Oligo.s4
SFG:Oligo.s0
SFG:Oligo.s1
SFG:Oligo.s2
SFG:Oligo.s3
Astro DCLK1
Astro GRM3
Astro CNTNAP2
Astro HSPH1
Astro ELMO1
Micro P2RY12
Micro NFIB
Micro HIF1A
Micro RPL*
Micro F13A1
OPC LHFPL3−AS1
OPC NEAT1 SYT1
Oligo SLC5A11 LRP2
Oligo RPLP1 UBB
Oligo SCD PMP22
Oligo CACNB2 FRMPD4
Endo NXN
Endo OST4
VLMC DCN
T−cell MGP
Fibro PDGFRB SLC19A1
Fibro PDGFRB SORBS1
Spearman correlation
Non−neuronal cells
1
0.5
0

**Supplementary Figures 3-5 | Cross-dataset correlation of cell type subpopulations.**

Heatmaps showing Spearman correlations of cell type subpopulations across independent datasets. Supplementary Figure 3 corresponds to the multi–brain region Allen Brain Institute dataset; Supplementary Figure 4 to the entorhinal cortex and superior temporal gyrus dataset from Leng et al. (2021); and Supplementary Figure 5 to the multiregional dataset from Mathys et al. (2024).

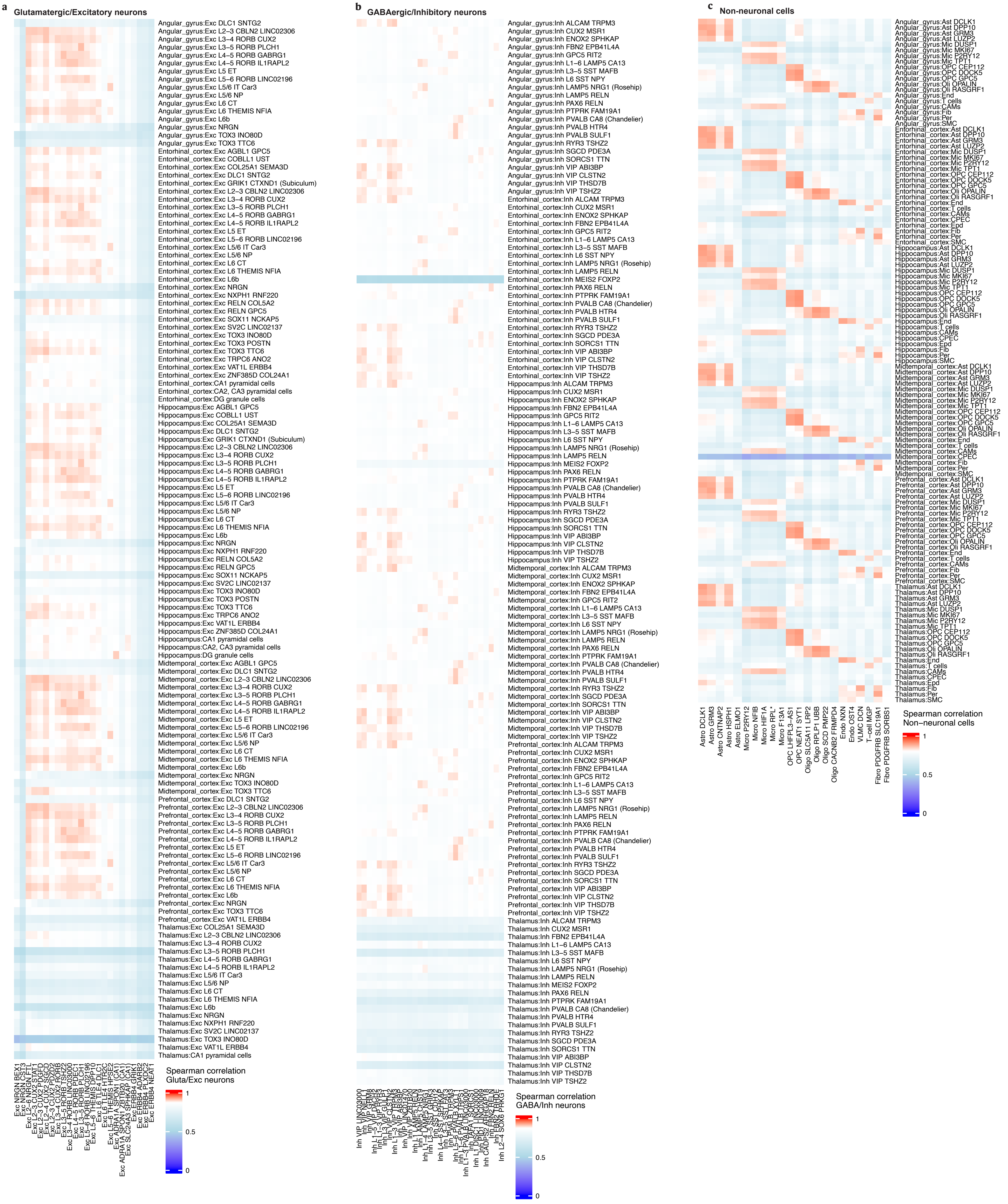
a
Glutamatergic/Excitatory neurons
Spearman correlation
Gluta/Exc neurons
1
0.5
0
b
GABAergic/Inhibitory neurons
Spearman correlation
GABA/Inh neurons
1
0.5
0
c
Non-neuronal cells
Spearman correlation
Non–neuronal cells
1
0.5
0

### Supplementary Figures 3-5 | Cross-dataset correlation of cell type subpopulations.

Heatmaps showing Spearman correlations of cell type subpopulations across independent datasets. Supplementary Figure 3 corresponds to the multi–brain region Allen Brain Institute dataset; Supplementary Figure 4 to the entorhinal cortex and superior temporal gyrus dataset from Leng et al. (2021); and Supplementary Figure 5 to the multiregional dataset from Mathys et al. (2024).
